# Post-intervention high-beta synchronization tracks stable-manifold alignment during stick-balancing learning

**DOI:** 10.64898/2026.10.27.762123

**Authors:** Haruki Shimokado, Kanato Hayashi, Taishin Nomura

## Abstract

Learning to stabilize an unstable object may depend not only on the accuracy of active corrective interventions but also on when to suspend intervention and allow the object’s intrinsic dynamics to evolve. We investigated learning-related changes in control dynamics and associated beta-band electroencephalographic (EEG) activity during virtual stick balancing. Ten healthy young adults practiced stabilizing a virtual inverted pendulum with a joystick over two consecutive days; scalp EEG was recorded from seven participants. For each control cycle, control quality was quantified as the distance (dist) between the pendulum state at the end of each active intervention phase and the stable manifold of the control-OFF subsystem. With practice, balance duration increased, dist decreased, and the proportion of time without active intervention increased. Time-frequency responses stratified by control quality showed that post-intervention high-beta event-related synchronization was most prominent in cycles with the smallest dist. Single-cycle linear mixed-effects models further revealed an inverse association between post-intervention high-beta power and dist after accounting for intervention magnitude and movement direction, as well as trial progress where applicable. This relationship was evident in both the initial Control and final Skilled task datasets. These findings indicate that stick-balancing practice improves the ability to terminate active intervention near the stable manifold and suggest that post-intervention high-beta synchronization tracks the quality of the transition from active correction to the intrinsic dynamics of the control-OFF subsystem.

## 1. Introduction

Maintaining an unstable object in an upright position, as in stick balancing, provides a useful paradigm for investigating how the central nervous system controls physical instability in the presence of sensory feedback delays and internal and external noise [1]. Such behavior has been explained by predictive control based on internal models [2,3] and by intermittent control, in which active corrective input is repeatedly engaged and disengaged [4–9]. Skilled performance is associated with longer periods without active intervention [5,7,8], and appropriately timed suspension of feedback control can improve the robustness of delayed unstable systems [10]. Thus, successful balancing may depend not only on how accurately corrective actions are generated, but also on when active correction is withheld—“knowing when not to control.”

A state-dependent intermittent-control model provides a dynamical explanation for this principle [11]. The model switches between a control-ON subsystem, in which delayed feedback torque is applied, and a control-OFF subsystem, in which the object evolves according to its intrinsic dynamics. Although the upright equilibrium of the control-OFF subsystem is unstable, it has a saddle structure with a stable manifold. When active intervention ends with the system state close to this manifold, the state transiently approaches the upright equilibrium without further corrective input.

Closer alignment with the stable manifold at intervention offset should therefore prolong the subsequent control-OFF period and reduce the need for continuous correction. Intermittent control models can reproduce characteristic nonlinear features of human balancing [4,5,8,12], can emerge through reinforcement learning [13], and have shown favorable model evidence relative to continuous feedback control in studies using Bayesian data assimilation [9,14]. However, how stable-manifold alignment changes during human motor learning, and whether its cycle-to-cycle variation is represented in cortical activity, remain unclear.

Beta-band activity is a candidate neural correlate of transitions between active intervention and its suspension. During quiet standing, centroparietal high-beta event-related synchronization (ERS) follows brief corrective motor activity [15] and accompanies periods of reduced active correction [16]. More generally, beta synchronization has been associated with maintenance of the current sensorimotor state [17,18], whereas post-movement beta activity has been linked to outcome processing, error evaluation, and confidence in internal-model estimates [19–24]. These observations raise the possibility that post-intervention beta synchronization is not merely a consequence of movement termination, but varies with the quality of the state achieved by the preceding intervention. In stick balancing, this possibility can be tested by relating beta power in each control cycle to the distance between the state at intervention offset and the stable manifold of the control-OFF subsystem. To our knowledge, no study has directly examined this relationship during learning.

Here, we recorded behavior and scalp electroencephalography (EEG) while healthy participants practiced a virtual stick-balancing task over two consecutive days. We quantified the quality of each control cycle by the stable-manifold distance at the end of active intervention (dist) and tested whether balance improvement was accompanied by reduced dist and increased time without active intervention. We then examined whether post-intervention high-beta activity covaried with dist at the single-cycle level while accounting for intervention magnitude, movement direction, and trial progress. We hypothesized that practice would improve stable-manifold alignment and that post-intervention high-beta synchronization would be stronger following interventions that placed the system closer to the stable manifold.

## 2. Methods

### 2.1 Participants

Ten healthy right-handed young adults (age range, 20–22 years; six men and four women) with no history of neurological disease or motor impairment participated in the study. All participants provided written informed consent before participation. Seven of the ten participants also consented to simultaneous electroencephalographic (EEG) recording. The experimental protocol was approved by the Ethics Committee of the Graduate School of Informatics, Kyoto University (approval no. KUIS-ERA-2025-009), and the study was conducted in accordance with the Declaration of Helsinki.

### 2.2 Virtual stick-balancing task and experimental protocol

Participants stabilized a virtual inverted pendulum displayed on a monitor by laterally tilting a joystick (T.16000M FCS, Thrustmaster, France) with both hands (Fig. 1A). They pushed the joystick leftward with the right hand and rightward with the left hand. All participants had normal or corrected-to-normal vision. The pendulum was modeled as a uniform rigid rod connected to the ground by a pin joint and governed by

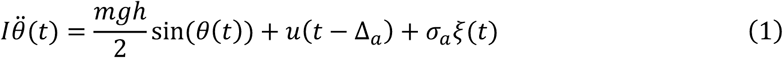

**Figure 1.**
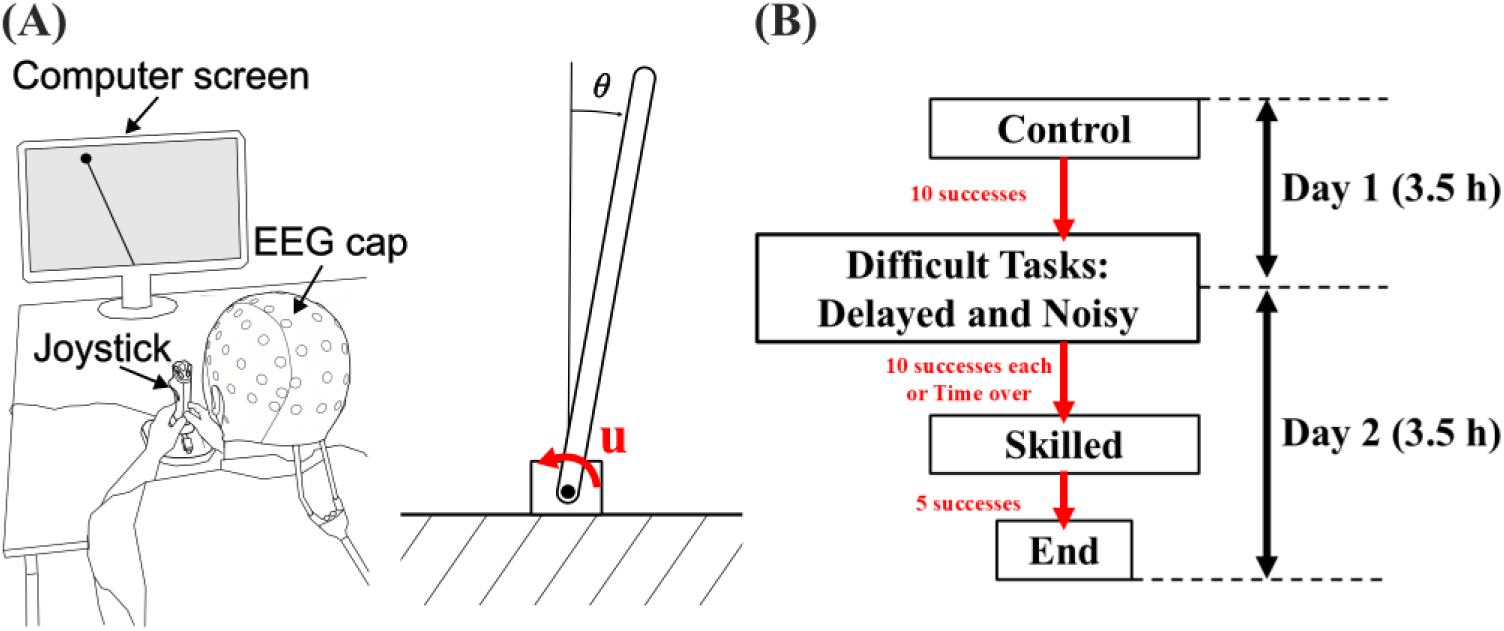
Virtual stick-balancing task and experimental protocol. (A) Schematic of the experimental setup. Participants manipulated a joystick with both hands in the horizontal plane to stabilize a virtual inverted pendulum displayed on a monitor while scalp electroencephalography was recorded. The pendulum dynamics were governed by Eq. (1), which describes the relationship among the pendulum angle *θ*, control torque *u*, artificial feedback delay, and process noise. (B) Two-day experimental protocol. Participants first performed the Control task and subsequently completed the two challenge tasks—the Delayed and Noisy tasks—in a randomized order. The Skilled task, which had the same environmental parameters as the Control task, was performed during the final stage of the experiment.

where *θ* is the pendulum angle, *I* = *m*ℎ^2^⁄3 ≈ 1.30 kgm^2^ is the moment of inertia, *g*=9.81 m⁄s^2^, *m*=0.625 kg, and ℎ=2.5 m. The normalized joystick output (−1 to 1) was multiplied by approximately 9.2 Nm (≈ 0.6 *mg*ℎ) to generate the control torque *u*. *ξ*(*t*) represents standard Gaussian white noise scaled by*σ_a_*. The dynamics were simulated in real time in MATLAB/Simulink (MathWorks, Natick, MA, USA) with a time step of 0.005 s using the forward Euler method, or the Euler–Maruyama method when process noise was applied.

Each trial lasted 130 s of cumulative task time. Participants attempted to maintain |*θ*| ≤ *π*/20 rad. Exceeding this range was counted as a fall, after which the pendulum was reset upright and the trial continued. Trials completed without a fall were classified as successful.

Task difficulty was manipulated by artificial control delay Δ*_a_* and process noise intensity *σ_a_*. Their values were (Δ*_a_*, *σ_a_*) = (0, 0) for the Control task, (0.07 s, 0) for the Delayed task, and (0, 0.05 Nm) for the Noisy task. The Delayed and Noisy tasks are collectively referred to as the challenge tasks.

The experiment was conducted over two consecutive days (Fig. 1B). Participants first performed the Control task until completing 50 trials or accumulating 10 successful trials, followed by the two challenge tasks in randomized order under the same stopping criteria. Testing was limited to 3.5 h per day; an unfinished task was resumed on Day 2. After the challenge tasks, participants repeated the Control condition until accumulating five successful trials; this final block was designated the Skilled task. If the challenge tasks were not completed, the Skilled task began after approximately 3 h of testing on Day 2. Pendulum angle, angular velocity, and control torque were recorded at 200 Hz.

### 2.3 Control-cycle identification and behavioral measures

Torque OFF and ON intervals were defined by |*u*| < 0.1 Nm and |*u*| ≥ 0.1 Nm, respectively. Torque onset and offset corresponded to OFF-to-ON and ON-to-OFF transitions. A control cycle extended from one torque onset to the next and comprised an ON interval followed by an OFF interval. For event-related analyses, an analysis epoch additionally included the immediately preceding OFF interval (pre-OFF interval); consequently, the OFF interval of one cycle also served as the pre-OFF interval of the next analysis epoch.

To exclude atypical interventions and operational artifacts, the pre-OFF, ON, and subsequent OFF intervals were separately normalized in time. For each trial and each torque direction, the mean time-normalized torque trajectory was calculated and correlated with each corresponding single-epoch trajectory. Epochs with a Pearson correlation coefficient *r* < 0.925 were excluded from behavioral and EEG analyses.

For each retained cycle, stable-manifold utilization was quantified from the pendulum state at torque offset. Linearizing Eq. (1) around the upright equilibrium with *u* = 0 yields eigenvalues ±*λ*, where

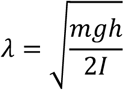

and the stable manifold is given by

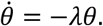

To place angle and angular velocity on a common scale, we used the normalized state plane (*θ*, *θ̇*/*λ*). The shortest Euclidean distance from the offset state to the stable manifold was defined as

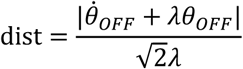

where *θ_OFF_* and *θ̇_OFF_* denote the angle and angular velocity at torque offset. Smaller dist indicated closer stable-manifold alignment and therefore higher control quality.

For each trial, we calculated the longest uninterrupted balance duration, mean dist across retained cycles, and proportion of the 130-s trial classified as OFF. Learning progression during the initial Control task was quantified as

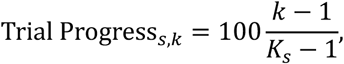

where *k* denotes the trial number and *K_s_* the total number of Control-task trials completed by participant *s*; the first and final trials therefore corresponded to 0% and 100%.

Control-task cycles at Trial Progress values of 0–25% and 75–100% were assigned to the Early and Late phases, respectively, and cycles from the final Skilled task to the Skilled phase. Independently of phase, retained cycles were classified by quartiles of the pooled dist distribution across the experiment. From the lowest to the highest quartile, the four control-quality levels were labeled Very Good, Good, Fine, and Bad.

### 2.4 EEG acquisition and processing

Scalp EEG was recorded in seven participants using a 64-channel mobile bioamplifier and a Waveguard cap (eego sports, ANT Neuro, Hengelo, the Netherlands). Ag/AgCl electrodes were positioned according to the international 10–10 system, with CPz as the online reference and AFz as ground. Electrode impedances were maintained below 25 kΩ [25], and signals were sampled at 2048 Hz.

Preprocessing was performed using EEGLAB v2022.0 [26]. The PREP pipeline [27] was used to attenuate 60-Hz line noise, interpolate noisy channels using spherical splines, and re-reference the data to a robust average. Signals were then band-pass-filtered at 1-70 Hz using a zero-phase, fourth-order Butterworth filter and downsampled to 200 Hz. The 1-Hz high-pass cutoff was used to improve stationarity before independent component analysis (ICA) [28].

Extended Infomax ICA was performed after principal component analysis, excluding components with covariance-matrix eigenvalues below 10^−7^. Independent components (ICs) were retained only when “Brain” had the highest ICLabel probability [29] and the fitted equivalent current dipole was located within the brain volume, defined as a depth value of ≤ 2 mm using ft_sourcedepth. No residual variance threshold was applied. Rejected ICs were visually inspected before removal, and artifact-reduced channel signals were reconstructed from the retained ICs. The number of retained brain ICs was 12.7 ± 3.7 per participant (mean ± SD).

### 2.5 Time-frequency analysis and cycle normalization

Time–frequency power *P*(*t*, *f*) was calculated for each EEG channel from 6 to 50 Hz using EEGLAB newtimef with the wavelet-cycle parameter [3 0.5], corresponding to a linear increase from 3 cycles at 6 Hz to 12.5 cycles at 50 Hz. Within each trial, power was normalized to its temporal mean at each frequency:

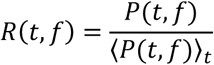

The normalized time-frequency response and corresponding pendulum-angle, angular-velocity, and torque signals were segmented into the analysis epochs defined in Section 2.3 and linearly time-warped separately for the pre-OFF, ON, and subsequent OFF intervals [30]. Across all participants and trials, the median durations of the OFF and ON intervals were 0.340 s (*mean* ± *SD* = 0.449 ± 0.351 s) and 0.485 s (0.575 ± 0.183 s), respectively. Based on their median durations, the normalized time axis assigned the preceding torque offset to −41%, torque onset to 0%, torque offset to 59%, and the next torque onset to 100%. Thus, the control cycle extended from 0% to 100%, with negative values representing the pre-OFF interval.

For descriptive event-related spectral perturbation (ERSP) maps, responses were averaged separately for each learning phase and control-quality level. To equalize participant contributions, the number of cycles was set to the minimum available across participants within each phase-by-quality combination, and that number was sampled once without replacement from each participant. Grand-average power was expressed in decibels as

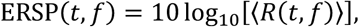

where the brackets denote averaging across the sampled cycles and participants. Subsampling was used only for descriptive ERSP maps; all retained cycles were included in the single-cycle statistical analyses.

### 2.6 Statistical analysis

#### 2.6.1 Behavioral analysis

Behavioral outcomes were analyzed using LMMs fitted with restricted maximum likelihood in MATLAB fitlme, with participant as a random intercept. For the initial Control task, separate models tested the fixed effect of Trial Progress on maximum uninterrupted balancing duration and mean dist. The proportion of time spent in the control-OFF state was compared between the Control and Skilled tasks using task condition as a fixed effect. Maximum uninterrupted balancing duration and mean dist were similarly compared among the Skilled, Delayed, and Noisy tasks, with Bonferroni correction applied to pairwise comparisons. Fixed-effect coefficients were evaluated using two-sided *t*-tests.

#### 2.6.2 Single-cycle EEG analysis

To examine cycle-level associations between EEG activity and behavioral variables, LMMs were fitted separately at each time–frequency pixel for each task and electrode (Cz and C3). For participant *s*, epoch *k*, time point *i*, and frequency bin *j*, the response was *y_sk_*(*i*, *j*) = *z*[*R_sk_*(*i*, *j*)]. The response and all continuous predictors were z-standardized within each model-fitting dataset. Intervention direction was coded as 0 for leftward and 1 for rightward interventions and was not standardized; interaction terms were constructed after standardization of the continuous predictors.

For the Skilled task, the fixed effects were dist, peak absolute control torque during the ON interval, direction, and the direction-by-peak torque interaction:

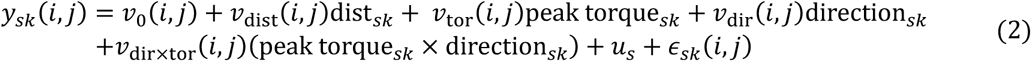

For the Control task, Trial Progress and its interaction with dist were additionally included:

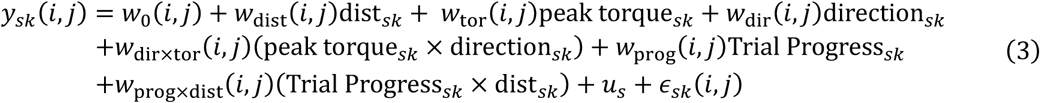

All coefficients were estimated separately at each time-frequency pixel; *u_s_* denotes the participant-specific random intercept and *ε_sk_*(*i*, *j*) the residual. Continuous main-effect coefficients represent standardized partial associations. Direction coefficients represent adjusted differences between rightward and leftward interventions, and *w*_prog×dist_ tests whether the association between dist and EEG power varied with Trial Progress.

Fixed effects were evaluated using two-sided *t*-tests. Benjamini–Hochberg false-discovery-rate correction [31] was applied separately for each task, electrode, and fixed effect across all time-frequency pixels. An FDR-adjusted *p*<0.05 was considered significant.

## 3 Results

### 3.1 Practice improves balancing performance and stable-manifold alignment while preserving intermittent control

Representative trajectories from the Control task showed that the pendulum state at torque offset was closer to the stable manifold in the Late than in the Early phase, allowing the control-OFF dynamics to bring the pendulum transiently toward the upright equilibrium (Fig. 2A).

**Figure 2.**
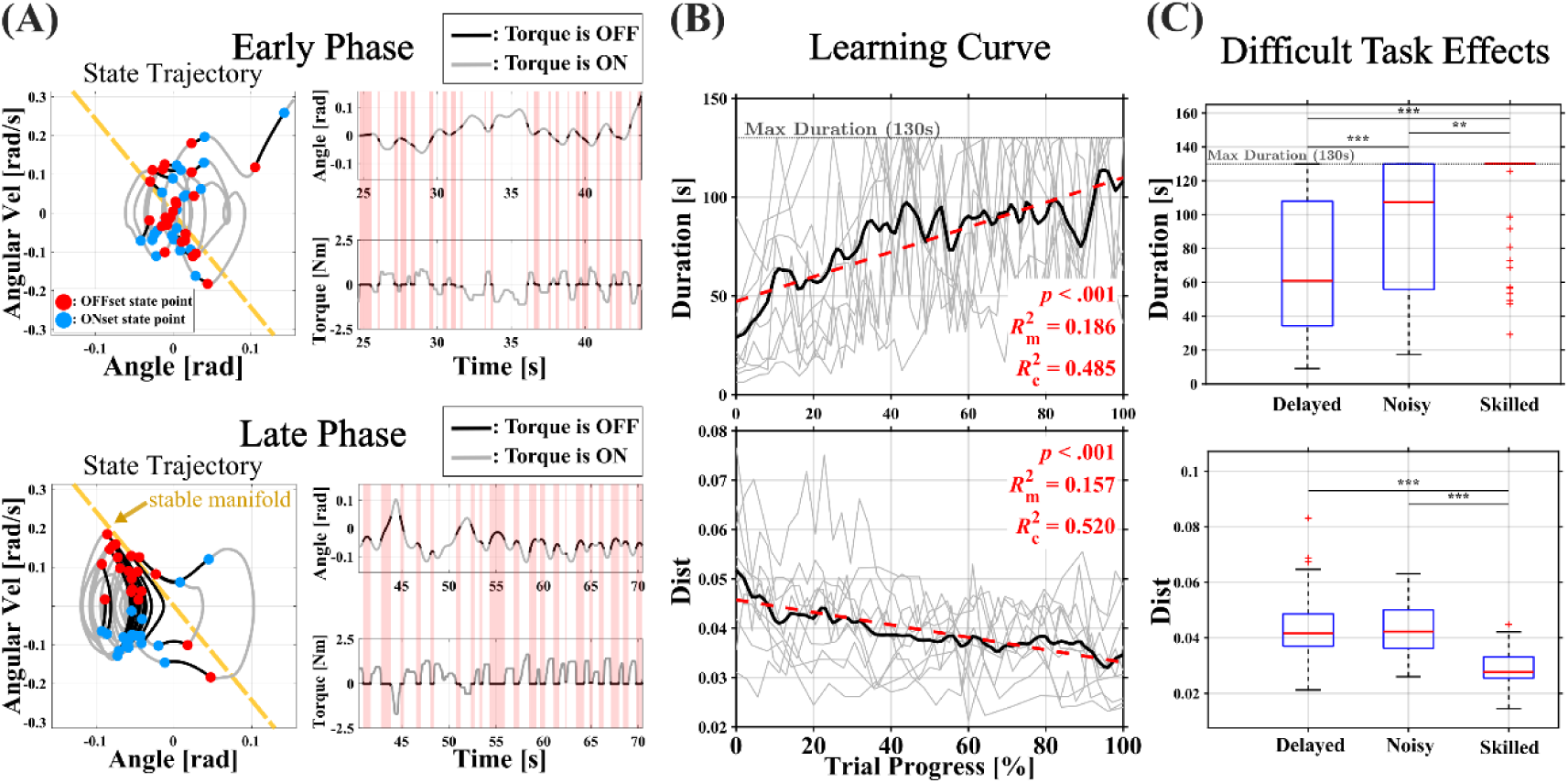
Practice-related improvements in balancing performance and stable-manifold alignment. (A) Representative state-plane trajectories in the angle–angular velocity plane and corresponding time series of pendulum angle, control state (ON or OFF), and control torque during the Early (top) and Late (bottom) phases of the Control task. Red and blue circles indicate the pendulum states at control-torque offset and onset, respectively. The yellow dashed line represents the stable manifold of the control-OFF subsystem. (B) Changes in the maximum uninterrupted balancing duration within each trial (top) and mean dist (bottom) as functions of Trial Progress (0–100%). Thin gray lines represent individual participants, thick black lines represent the across-participant averages, and red dashed lines indicate the fitted linear mixed-effects model (LMM). 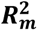 and 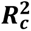 denote the marginal and conditional coefficients of determination, respectively. (C) Boxplots comparing the maximum uninterrupted balancing duration (top) and mean dist (bottom) among the Delayed, Noisy, and Skilled tasks. Reported significance levels are based on Bonferroni-adjusted comparisons. ^∗∗^*p*_adj_ < 0.01, ^∗∗∗^*p*_adj_ < 0.001.

Across Control-task trials, maximum uninterrupted balancing duration increased with Trial Progress (estimate = 0.63, *SE* = 0.050, *t*(df = 373) = 12.42, *p* < 0.001), whereas mean dist decreased (estimate = −1.3 × 10^−4^, *SE* = 1.1 × 10^−5^, *t*(df = 373) = −11.48, *p* < 0.001) (Fig. 2B). The same patterns were found among the seven EEG participants for balancing duration (estimate = 0.61, *SE* = 0.063, *t*(df = 241) = 9.76, *p* < 0.001) and mean dist (estimate = −1.4 × 10^−4^, *SE* = 1.4 × 10^−5^, *t*(df = 241) = −9.93, *p* < 0.001). The proportion of trial time classified as control-OFF was also greater in the Skilled task than in the initial Control task (estimate = 0.10, *SE* = 0.007, *t*(df = 427) = 14.57, *p* < 0.001), indicating that intermittent control persisted after learning.

Relative to the Skilled task, maximum uninterrupted balancing duration was shorter in both the Delayed (estimate = −49.25, *SE* = 4.73, *t*(df = 549) = −10.42, *p_adj_* < 0.001) and Noisy tasks (estimate = −19.47, *SE* = 5.66, *t*(df = 549) = −3.44, *p_adj_* = 0.002) (Fig. 2C). Mean dist was correspondingly greater in both the Delayed (estimate = 0.013, *SE* = 0.0010, *t*(df = 549) = 13.71, *p_adj_* < 0.001) and Noisy tasks (estimate = 0.012, *SE* = 0.0012, *t*(df = 549) = 10.56, *p_adj_* < 0.001). Balancing duration was longer in the Noisy than in the Delayed task (estimate = 29.79, SE = 3.99, *t*(549) = 7.46, *p_adj_* < 0.001), whereas mean dist did not differ between them (estimate=-0.0010, *SE* = 0.0008, *t*(549) = -1.23, *p_adj_* = 0.658).

Six of ten participants reached 10 successful trials during the initial Control task before the 50-trial limit, whereas all participants reached five successful trials in the final Skilled task. Because success required uninterrupted balancing for 130 s, Skilled-task duration showed a pronounced ceiling effect, supporting dist as a complementary cycle-level measure of control quality.

The dist quartiles defining Very Good, Good, Fine, and Bad levels were 0.0075, 0.0163, and 0.0294. Consistent with the decrease in mean dist, the proportion of Very Good cycles increased across the Early, Late, and Skilled phases and was highest in the Skilled phase (Table 1).

**Table 1.** Proportions of control cycles at each dist-based control-quality level across learning phases. Values are mean ± SD across participants.

|  | Very Good | Good | Fine | Bad |
| --- | --- | --- | --- | --- |
| **Early** | **20.1 ± 5.7%** | **22.5 ± 5.3%** | **23.8 ± 4.2%** | **33.6 ± 14.5%** |
| **Late** | **29.4 ± 6.2%** | **26.5 ± 5.0%** | **24.5 ± 1.8%** | **19.6 ± 9.6%** |
| **Skilled** | **37.7 ± 9.2%** | **32.7 ± 2.5%** | **20.4 ± 3.8%** | **9.3 ± 6.5%** |

### 3.2 Post-intervention high-beta activity is associated with stable-manifold alignment across individual control cycles

Figure 3 shows the descriptive ERSP at Cz across learning phases and control-quality levels. Time-normalized torque profiles were broadly similar across learning phases, whereas during the post-intervention OFF interval, the angle and angular velocity profiles of Very Good cycles approached the upright equilibrium more closely than those of lower-quality cycles.

**Figure 3.**
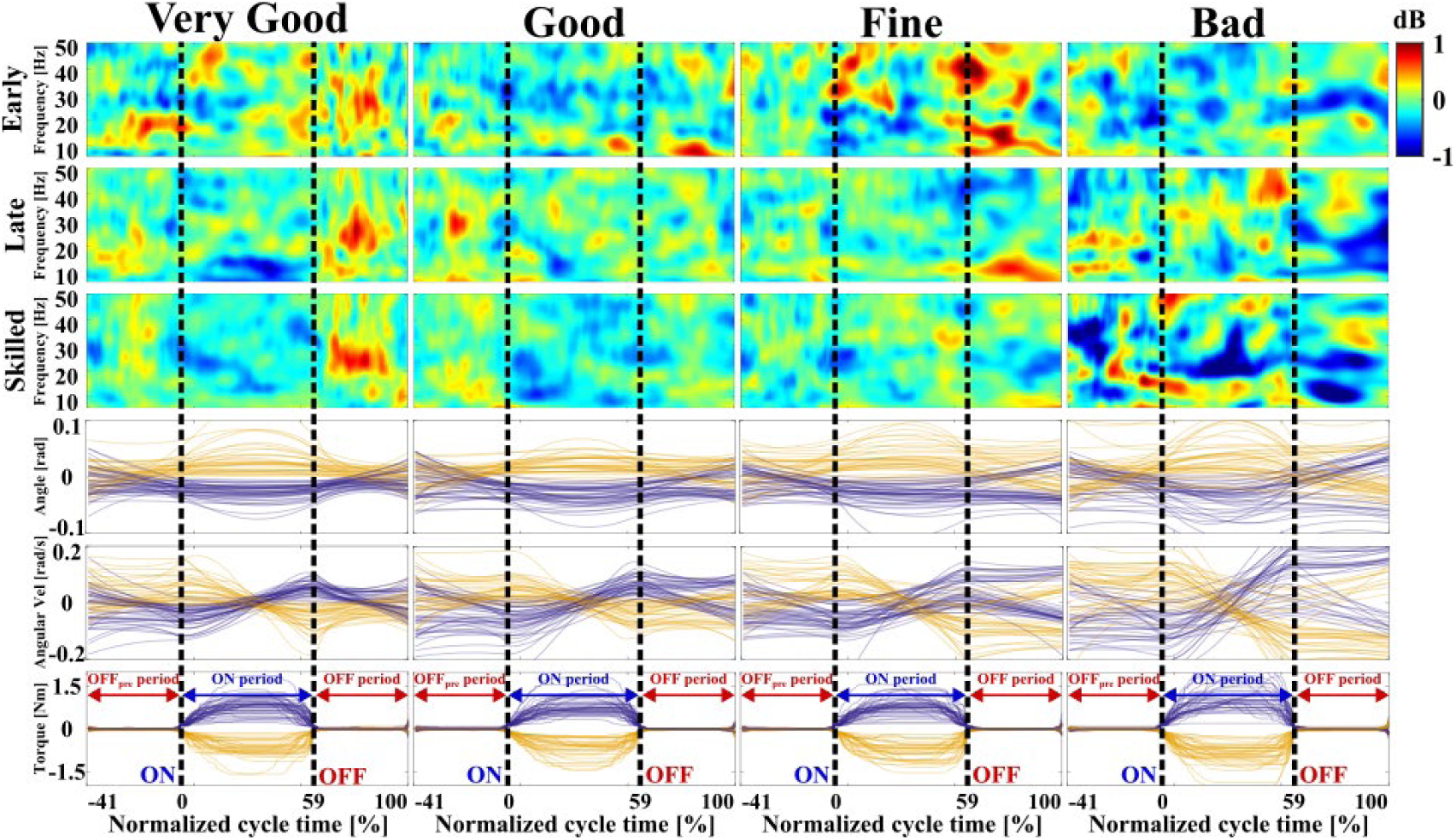
Post-intervention high-beta ERS across control-quality levels and learning phases. Grand-average event-related spectral perturbations (ERSPs) recorded at Cz for Very Good, Good, Fine, and Bad control cycles during the Early, Late, and Skilled phases (*N* = **77**). To equalize the contribution of each participant to each grand-average ERSP, the number of cycles was set to the smallest number available across participants within each combination of learning phase and control-quality level. The resulting numbers of cycles sampled without replacement from each participant were 62, 78, 87, and 85 for the Very Good, Good, Fine, and Bad levels, respectively, during the Early phase; 100, 91, 87, and 43 during the Late phase; and 150, 135, 76, and 22 during the Skilled phase. ERSPs are expressed in decibels. The analysis epochs were time-warped to a normalized time axis extending from −41% to 100%, comprising the pre-OFF interval (−41% to 0%), ON interval (0% to 59%), and subsequent OFF interval (59% to 100%). The black vertical dashed lines indicate control-torque onset (0%) and offset (59%). The lower three rows show the corresponding time courses of pendulum angle, angular velocity, and control torque. For visualization, 10 cycles per participant from the Skilled-task dataset are superimposed for each control-quality level. Purple and yellow traces distinguish the two joystick-intervention directions.

Across phases and control-quality levels, high-beta power at approximately 20–30 Hz decreased shortly before torque onset and during the ON interval. Following torque offset, post-intervention high-beta event-related synchronization (PI-high-β-ERS) was most apparent in Very Good cycles throughout the Early, Late, and Skilled phases. Within the Skilled phase, PI-high-β-ERS remained prominent in Very Good cycles but was less apparent at lower control-quality levels. Thus, the descriptive maps suggested that PI-high-β-ERS depended primarily on individual-cycle quality rather than increasing uniformly across all cycles with learning.

Single-cycle LMMs included 4,971 Skilled-task cycles and 20,667 Control-task cycles (Fig. 4). Variance inflation factors were ≤2.09, indicating no substantial multicollinearity. In both tasks, dist was negatively associated with post-intervention beta power at Cz and C3 during the OFF interval after adjustment for peak torque, direction, and their interaction. These associations survived FDR correction and broadly overlapped with the PI-high-β-ERS observed descriptively in Very Good cycles. Because smaller dist indicates closer stable-manifold alignment, the negative coefficients indicate greater post-intervention beta synchronization following higher-quality interventions. At C3, the association extended into lower beta frequencies.

**Figure 4.**
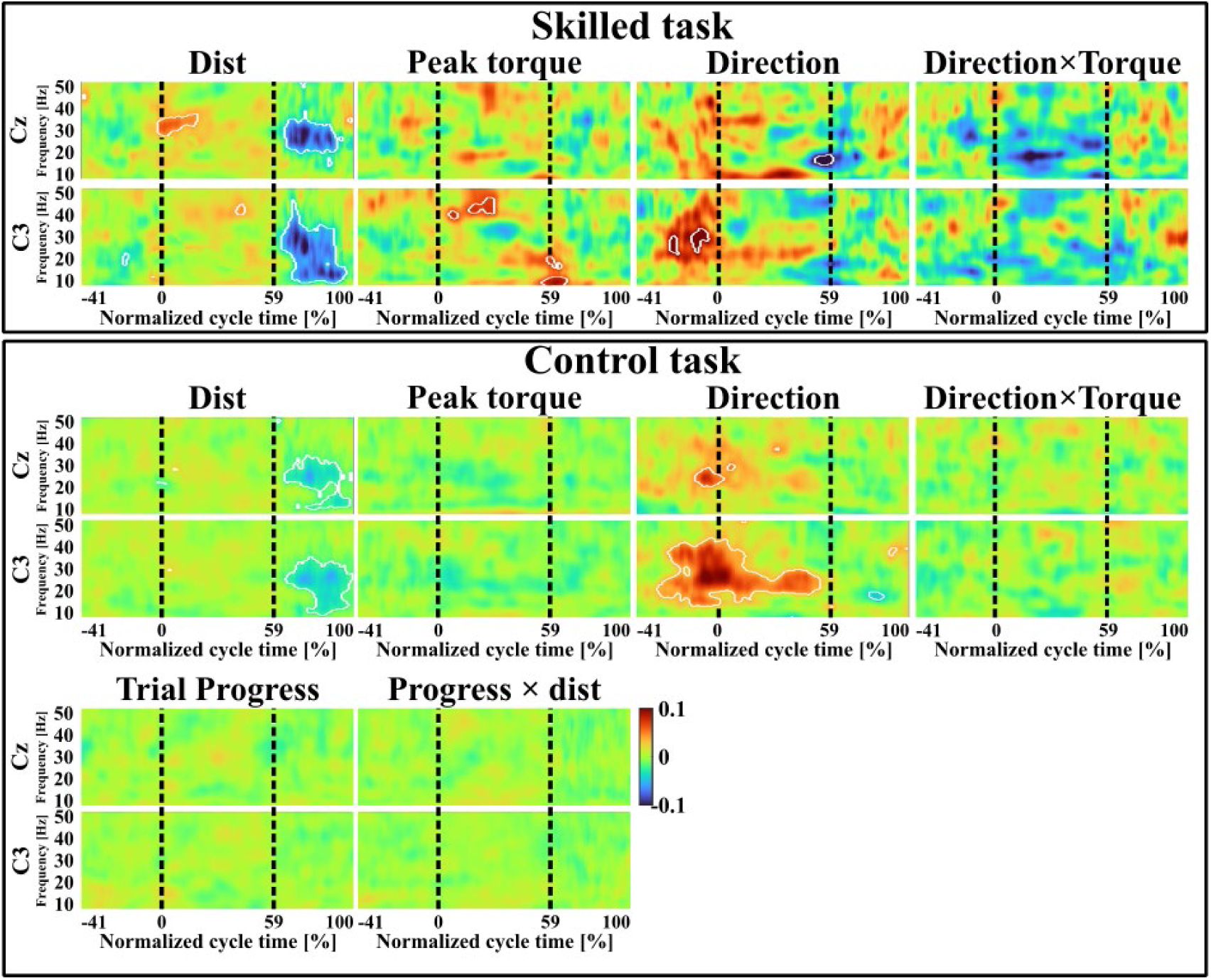
Time–frequency maps of single-cycle associations estimated using linear mixed-effects models. Maps of the fixed-effect coefficients from the linear mixed-effects models fitted to the normalized time–frequency EEG power at Cz and C3. The upper section shows the coefficients from the Skilled-task model in Eq. (2): dist (*v*_dist_), peak control torque (*v*_tor_), intervention direction (*v*_dir_), and the direction-by-torque interaction (*v*_dir×tor_). The lower section shows the coefficients from the Control-task model in Eq. (3): dist (*w*_dist_), peak control torque (*w*_tor_), intervention direction (*w*_dir_), the direction-by-torque interaction (*w*_dir×tor_), Trial Progress (*w*_prog_), and the Trial Progress-by-dist interaction (*w*_prog×dist_). Intervention direction was dummy-coded as 0 for leftward and 1 for rightward interventions. White contours enclose time–frequency regions that remained significant after Benjamini–Hochberg false-discovery-rate correction (FDR-adjusted *p* < 0.05). Black vertical dashed lines indicate control-torque onset (0%) and offset (59%). Positive and negative coefficient values indicate positive and negative adjusted associations with normalized EEG power, respectively.

In the Skilled task, dist was also positively associated with power near 30 Hz around torque onset; thus, cycles with smaller dist at the subsequent torque offset exhibited a greater reduction in high-beta power at intervention onset.

Additional covariate effects occurred in restricted regions. Peak torque was associated with C3 power primarily during the Skilled-task ON interval, whereas direction effects occurred around torque onset, particularly at C3. These coefficients represent adjusted differences between rightward and leftward interventions and are distinct from the association between dist and PI-high-β-ERS.

Neither Trial Progress nor its interaction with dist showed an FDR-significant region in the Control task. Thus, the cycle-level association between dist and post-intervention beta power was detectable during initial and skilled performance, without evidence that its magnitude changed systematically over the initial Control task.

## 4 Discussion

### 4.1 Principal findings

This study identified two principal findings during practice of a virtual stick-balancing task. First, balancing duration increased and the pendulum state at intervention offset approached the stable manifold, while the proportion of time spent in the control-OFF state increased. Thus, performance improved while intermittent control was preserved. Second, post-intervention beta power was negatively associated with dist at the individual-cycle level in both the Control and Skilled tasks after adjustment for intervention magnitude, direction, and learning progress where applicable; this relationship was also descriptively apparent across learning phases. Together, these findings suggest that practice improves the termination of active intervention near a dynamically advantageous state and that post-intervention high-beta activity reflects the quality of the resulting transition into control-OFF dynamics.

### 4.2 Practice refines when active intervention is terminated

A defining feature of intermittent control is that stabilization is achieved not by continuously correcting deviations but by alternating active intervention with periods governed by the intrinsic dynamics of the controlled system [4–9,11]. Because the control-OFF subsystem has a saddle-type equilibrium, a state placed near its stable manifold transiently approaches the upright equilibrium without further corrective torque. Dist therefore quantifies how favorably active intervention terminates.

The increase in balancing duration and decrease in dist indicate that practice improved this termination. However, ON–OFF control was present throughout learning: high-quality cycles occurred during the Early phase, whereas poorly aligned cycles remained during the Skilled phase. Rather than reflecting replacement of one control mode by another, this pattern suggests increased reliability in terminating interventions near the stable manifold. The present data cannot distinguish gradual adjustment of control parameters from more frequent selection of previously discovered high-quality actions. Because no trial-by-trial policy-update model was fitted, reinforcement learning, bandit-like exploration, and softmax policy updating remain hypotheses requiring explicit model comparison.

Terminating intervention near the stable manifold may both exploit the convergent component of the control-OFF dynamics and reduce delayed overcorrection [10,11]. It may also reduce the information required to map continuously varying sensory states onto corrective actions, consistent with policy complexity defined by state–action mutual information [32]. However, policy complexity was not quantified here, and its relationship to stable-manifold utilization remains to be tested computationally.

### 4.3 Functional interpretation of post-intervention high-beta synchronization

Post-movement beta synchronization has been linked to movement termination, maintenance of the current sensorimotor state, outcome processing, and confidence in internal-model estimates [17–24]. The present findings extend these observations to repeated interventions during ongoing unstable control: PI-high-β-ERS was strongest after favorable stable-manifold alignment, and post-intervention beta power was negatively associated with dist in both tasks after adjustment for intervention magnitude and direction.

PI-high-β-ERS may therefore mark temporary disengagement of feedback-driven corrective action after the system enters a favorable dynamical state. This interpretation does not imply interruption of sensory input or functionally open-loop control; sensory information may remain monitored without being immediately transformed into corrective torque. This view is consistent with prolonged beta synchronization following small postural recoveries and its proposed relationship to active monitoring [33]. Nevertheless, the present data cannot distinguish suppression of further correction from monitoring or evaluation of the post-intervention state.

In the Skilled task, smaller dist at the subsequent torque offset was also associated with a greater reduction in high-beta power near intervention onset. This earlier activity may reflect a preparatory or intervention-related state associated with a favorable outcome, although the association does not establish causality. Direction effects at C3 may instead reflect lateralized movement preparation or different hand contributions [34] and should be distinguished from the post-intervention association with dist.

A cortico-basal ganglia-thalamic network model has shown that beta dynamics can accompany switching between active and inactive control states [35], providing a possible circuit-level framework for these findings. However, scalp EEG cannot establish basal-ganglia involvement, including hyperdirect-pathway or STN–GPe interactions, dopaminergic evaluation, or internal reward. Nor does the C3 low-beta association establish that cortical high-beta activity generated a secondary basal-ganglia response. These mechanisms remain testable hypotheses rather than conclusions of this study.

### 4.4 Limitations and implications

EEG was analyzed in seven participants. Although the large number of control cycles improved within-participant precision, it does not substitute for a larger number of independent participants or ensure population-level generalizability. The random-intercept LMMs accounted for clustering within participants, but the sample was insufficient to estimate more complex between-participant variation reliably. In addition, dist is derived from the linearized control-OFF dynamics and captures only one aspect of control quality, although its decrease paralleled improved balancing performance.

Time warping and wavelet smoothing may have attenuated or displaced brief temporal features, limiting precise claims about the temporal ordering of neural and behavioral events. Moreover, leftward and rightward interventions involved different hand contributions, preventing direction effects from being separated fully from effector laterality.

Despite these limitations, the convergence of improved performance, reduced dist, and cycle-resolved beta activity supports a restrained but mechanistically informative conclusion: practice improves the termination of active intervention near the stable manifold, while post-intervention high-beta synchronization reflects the quality of the resulting transition into control-OFF dynamics. This framework provides a testable basis for investigating how the nervous system learns not only how to intervene, but also when further intervention is unnecessary.

## Data and code availability

Data and analysis code supporting the findings of this study are available from the corresponding authors upon reasonable request.

## Acknowledgements

This work was supported by JSPS KAKENHI Grant Number JP22H03662 (T.N.).

## Declaration of competing interest

The authors declare no competing interests.

## Notes

### Competing Interest Statement

The authors have declared no competing interest.

